# Stage-specific extracellular vesicle cargo from Schwann cells orchestrates peripheral nerve regeneration

**DOI:** 10.1101/2025.09.08.674933

**Authors:** Manju Sharma, Supasek Kongsomros, Maulee Sheth, Somchai Chutipongtanate, Leyla Esfandiari

## Abstract

Schwann cells (SCs) play a critical role in peripheral nerve regeneration, undergoing dynamic phenotype transitioning from myelinating to repair stages following injury. While SC-derived extracellular vesicles (SC-EVs) have emerged as key mediators of intercellular communication during nerve repair, their stage-specific molecular cargo and functional roles remained incomplete understood. Here, we delineate protein, microRNA and lncRNA landscapes of SC-EVs across distinct differentiation stages, including immature, myelinating, and repair phenotypes, using an *in vitro* model of primary rat SCs. We show that myelinating SC-EVs are enriched with reprogramming factor SOX2 and neurotrophin receptor p75^NTR^, while repair SC-EVs carry distinct microRNAs predicted to modulate genes involved in myelin ensheathment, neuronal differentiation and neurogenesis. Moreover, repair SC-EVs contain long non-coding RNAs (lncRNAs) that may regulate miRNA activity. These findings reveal a novel mechanism by which SC-EVs orchestrate neuronal regeneration through stage-specific molecular cargo, and establishes a foundational model for investigating SC plasticity in peripheral nerve repair.

## Introduction

Schwann cells (SCs), the glial cells of the peripheral nervous system (PNS), play a fundamental role in maintaining neuronal structure and function (1). These cells form the myelin sheath, which wraps around the axon and enables saltatory conduction of nerve impulses at the nodes of Ranvier, thereby facilitating rapid signal transmission (2). During development, immature SCs undergo a morphogenetic process called radial sorting, whereby immature SCs segregate large-caliber axons (>1 µm) from mixed bundles in response to signals from both basal lamina and axons (3). This process directs SC differentiation into myelinating SCs, which ensheathe single large-caliber axons in a 1:1 ratio, or non-myelinating SCs, which surround multiple small-caliber axons to form Remak bundles (3).

Schwann cells also play a critical role in peripheral nerve repair following injury (4, 5). Axonal damage triggers the activation of transcription factors such as c-Jun, STAT-3, and Mitf, which reprogram myelinating SCs into a repair phenotype (6–8). Elevated c-Jun drives myelin clearance via JNK/c-Jun-dependent myelinophagy (8) and concurrently suppresses the myelination-promoting transcription factor KROX20/Egr2 to maintain SCs in a repair stage (9). In parallel, repair SCs facilitate debris clearance by recruiting macrophages through TNF-α and MCP-1 (10) and guide the axonal regrowth by aligning into bunger bands, which are regulated by Ephrin-B2/EphB2 signaling and Sox2-mediated N-cadherin redistribution (11). To promote axon elongation and neuronal survival, repair SCs secrete neurotrophic factors, including glial cell line-derived neurotrophic factor (GDNF), neurotrophin-3 (NT3), brain-derived neurotrophic factor (BDNF), and nerve growth factor (NGF) (5, 11, 12). Upon completion of axonal regrowth, neuregulin-1 type III (Nrg1 III) activates ErbB2/B3 receptors on repair SCs, initiating PI3K/Akt and MAPK/Erk1/2 signaling cascades that restore KROX20 and SOX10 expression and enable remyelination (13).

Most current research on SC-mediated repair has focused on the direct role of cells and cell-cell interactions. However, growing evidence suggests that extracellular vesicles (EVs) also play a critical role in facilitating SC-neuron communication during peripheral nerve regeneration (14). EVs are nanoscale, lipid bilayer-enclosed particles released by cells that transport bioactive molecules, including proteins, lipids, mRNAs, miRNAs, and lncRNAs to mediate intercellular communication over short and long distances (15, 16). EVs also regulate immune responses, promote tissue repair, and modulate disease progression (17, 18). In the context of nerve regeneration, EVs from various sources, including mesenchymal stem cells (MSCs), adipose-derived stem cells (ASCs), neurons, olfactory ensheathing cells, and dental pulp stem cells, have been shown to promote axonal growth **and myelination** (**19–22**). Schwann cell-derived extracellular vesicles (SC-EVs), in particular, exhibit potent neuroprotective and regenerative properties (23–26). SC-EVs enhance retinal ganglion cell survival and axonal growth following optic nerve injury (27) as well as to promote motoneuron regeneration (28). Mechanistically, SC-EVs deliver regeneration-associated molecules, such as miR-21 (enriched in repair SCs) and miR-23b-3p, both of which promote neurite outgrowth (26, 29). Additionally, SC-EVs carry TNFR1, which binds and sequesters excess TNFα, thereby limiting inflammation and promoting macrophage polarization toward a pro-repair phenotype, ultimately supporting tissue regeneration (30). These findings emphasize the regenerative potential of SC-EVs.

Despite promising evidence, the functional role of SC-EVs in nerve repair remains incomplete understood, particularly regarding their contributions at distinct stages of SC differentiation. We hypothesize that SC-EVs, whether released from repair SCs at injury sites or from adjacent uninjured myelinating SCs, act as mediators of nerve regeneration by delivering stage-specific molecular cargo to neurons and SCs, thereby coordinating the repair process. In this study, we investigated the roles of SC-EVs across three differentiation stages: immature, myelinating, and repair SCs. By isolating and characterizing EVs from each phenotype, we performed western blotting and comparative transcriptomic profiling to identify regeneration-associated proteins and RNAs. Our findings reveal how SC-EVs contribute to neuronal repair and SC plasticity, providing mechanistic insights into SC-EV-mediated regeneration and enhancing our understanding of EV-mediated intercellular signaling in peripheral nerve repair.

## Results

### 1. Validation and characterization of an *in vitro* Schwann cell differentiation model

Schwann cells (SCs) are essential for peripheral nerve repair, undergoing reprogramming from a myelinating to a repair phenotype in response to injury. To investigate the molecular mechanisms underlying this transition and EV-mediated communication, we utilized a previously established *in vitro* differentiation model (**Figure 1A**) that recapitulates key stages of SC maturation and reversion (31). Primary rat SCs were treated with dbcAMP to simulate axonal contact and activate signaling cascades that induce differentiation into the myelinating phenotype (32). Subsequent dbcAMP withdrawal prompted reversion to the repair phenotype.

**Figure 1.**
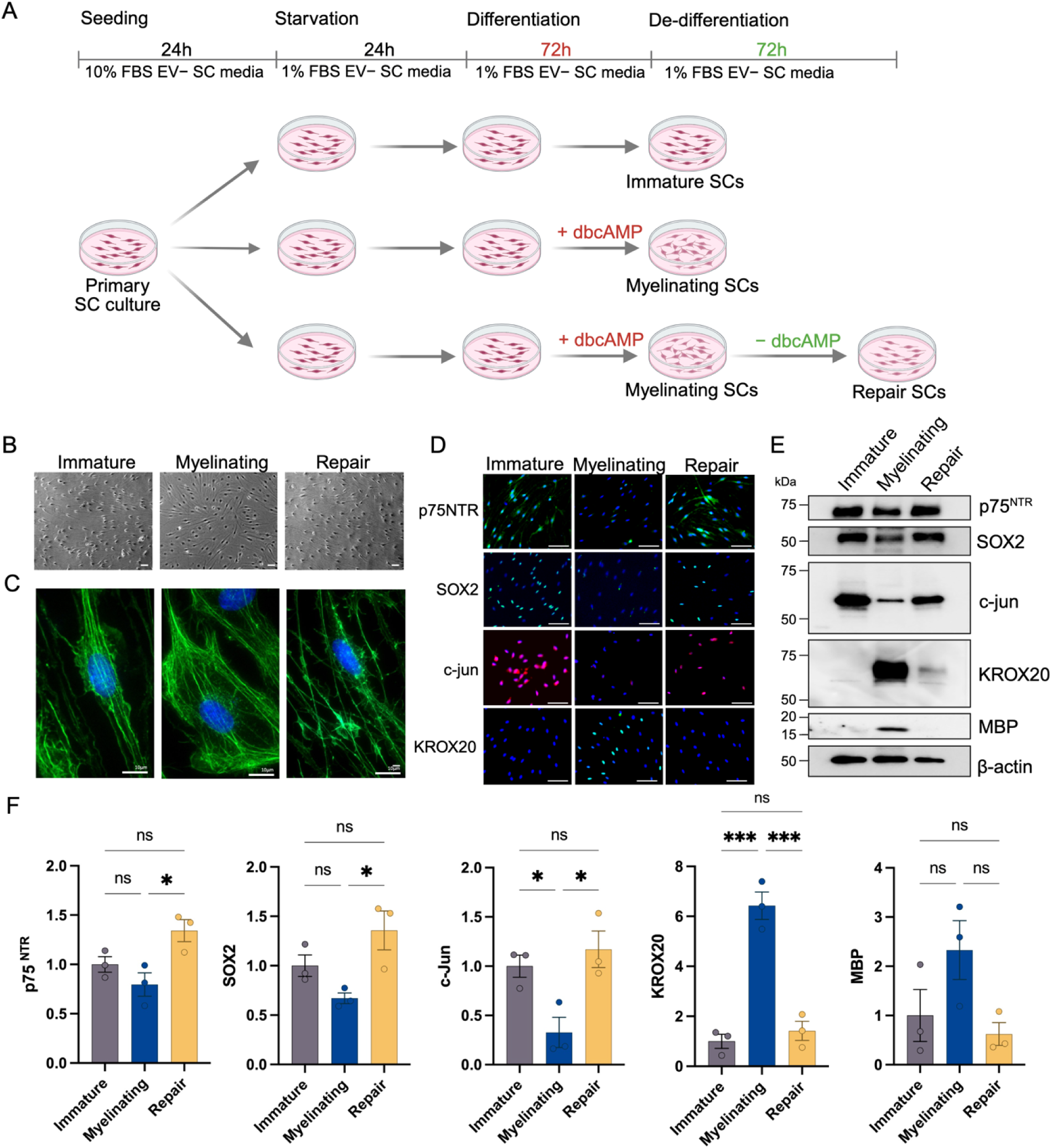
Schwann cell differentiation and molecular marker validation. (A) Schematic of the *in vitro* differentiation model used to induce Schwann cells (SCs) to induce myelination, followed by reversion to a repair phenotype. Rat primary SCs were treated with 1 mM dbcAMP in rat SC basal media for 72 h to induce differentiation into myelinating SCs, followed by dbcAMP withdrawal to reprogram into repair SCs. (B) Phase contrast and (C) phalloidin-stained super-resolution fluorescence imaging illustrates morphological differences and cytoskeletal organization in immature, myelinating, and repair SCs. Scale bars: 20 µm and 10 µm, respectively. (D) Immunofluorescence staining for SC-specific markers p75^NTR^, SOX2, c-Jun, and KROX20 shows distinct molecular profiles for differentiation stages. Immature SCs express high p75^NTR^, c-Jun, and Sox2; myelinating SCs express KROX20; repair SCs re-express immature proliferative markers. Scale bars: 100 µm. (E) Western blot analysis confirms different expression patterns of SC markers across distinct differentiation stages (the full-length blot images were available in **Supplementary** Figure 1). (F) Western blot quantification shows downregulation of p75^NTR^, c-Jun, and Sox2 as normalized to actin in myelinating SCs, with significant upregulation in repair SCs. KROX20 and MBP were elevated in myelinating SCs. The data are presented as the mean ± SEM of three biological replicates. Statistical analysis was performed by using one-way ANOVA with Tukey’s multiple comparisons test: * p < 0.05, ** p < 0.005, and *** p < 0.001, ns, not significant.

Distinct morphological changes of SC differentiation were observed under phase-contrast and actin-stained imaging (**Figure 1B, C**). Immature SCs exhibited an undifferentiated, spindle-shaped, elongated morphology, while myelinating SCs were bigger, rounded, and flattened appearance with reduced cytoskeletal organization, consistent with the previous report (32), indicating activation of the myelination process. Upon dbcAMP withdrawal, repair SCs reverted to an elongated morphology and re-established organized actin filaments, resembling the immature SC phenotype.

To further evaluate stage-specific characteristics, we assessed the expression of key markers by immunofluorescence (**Figure 1D**) and western blot analysis (**Figure 1E**). Immature SCs expressed transcription factors c-Jun and SOX2 and the neurotrophin receptor p75^NTR^, indicative of a proliferative, undifferentiation stage. Myelinating SCs showed upregulation of KROX20 and myelin basic protein (MBP), hallmarks of myelin sheath formation. In contrast, repair SCs upregulated c-Jun, SOX2, and p75^NTR^, and concurrently repressed KROX20, consistent with a regenerative, dedifferentiation stage. Quantification of protein expression confirmed significant upregulation of P75^NTR^, SOX2, and c-Jun expressions in repair SCs, and increased expression of KROX20 and MBP in myelinating SCs (**Figure 1F**). Together, these findings validate the *in vitro* SC differentiation model as a robust platform for dissecting molecular transitions and SC-EV-mediated communication during nerve injury and regeneration.

### 2. Transcriptomic profiling reveals molecular pathways driving Schwann cell myelination and repair

To elucidate molecular mechanisms governing SC differentiation, we performed poly(A) RNA sequencing on SCs at distinct differentiation stages; immature, myelinating, and repair phenotypes. The heatmap with hierarchical clustering showed discrete gene expression profiles among these states (Figure 2A). Principal component analysis (PCA) revealed clear transcriptional segregation between immature and myelinating SCs (Figure 2B), as well as between myelinating and repair SCs (Figure 2C). Notably, immature and repair SCs exhibited overlapping transcriptional signatures (Figure 2D), consistent with a partial reversion to an immature-like stage during nerve repair (1, 33).

**Figure 2.**
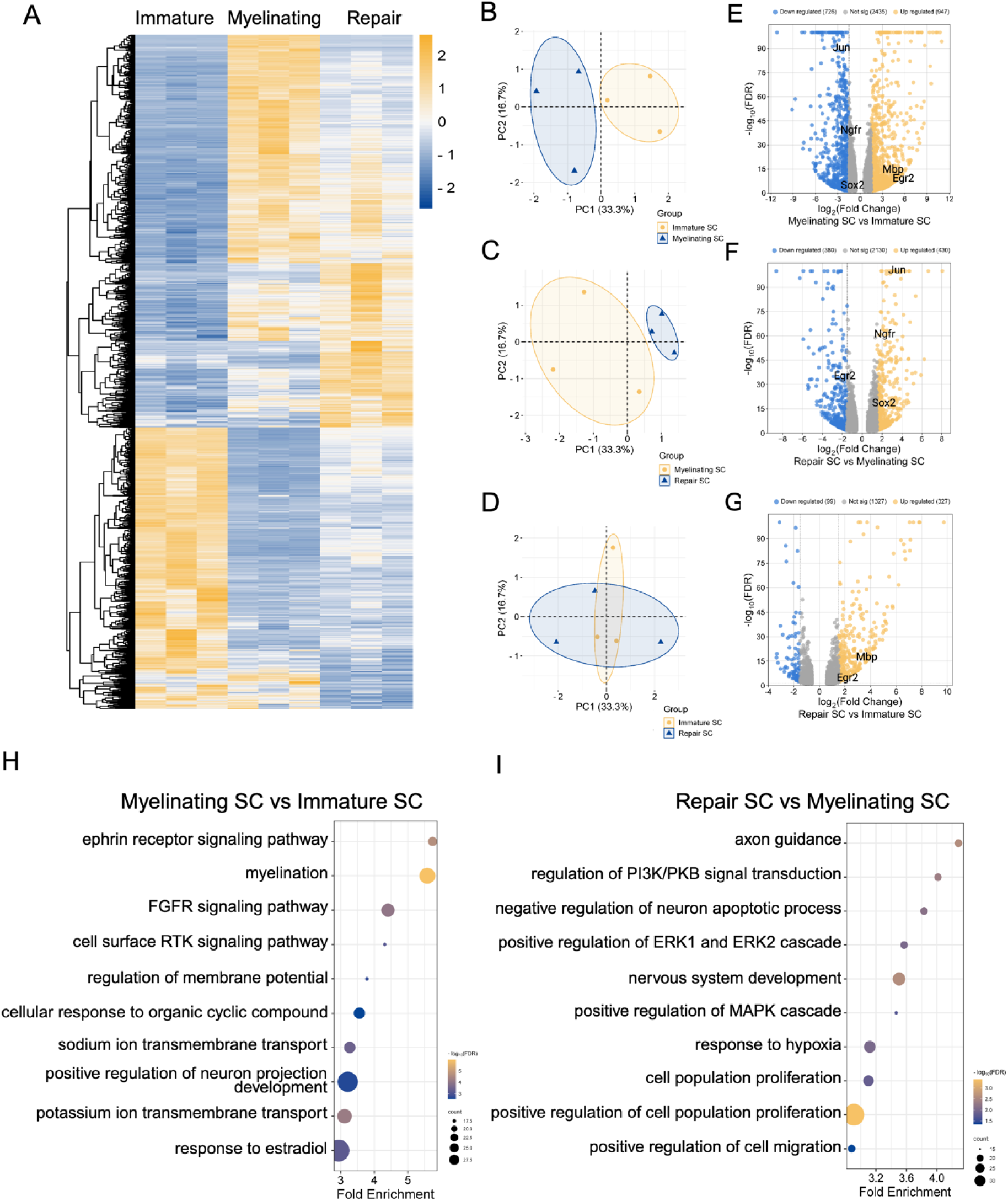
Transcriptomic analysis of Schwann cell differentiation. (A) Heatmap with hierarchical clustering shows significant differential gene expression profiles across immature, myelinating, and repair SCs. Each row represents an individual mRNA, with color intensity indicating expression levels (blue: downregulated; yellow: upregulated). Principal component analysis (PCA) reveals distinct clustering of transcriptomic profiles between myelinating and immature SCs (B), repair and myelinating SCs (C), and overlapping profiles between repair and immature SCs (D). (E-G) Volcano plots demonstrate differentially expressed genes between SC stages, with upregulated (yellow) and downregulated (blue) genes. Genes with log₂ fold change ≥ 1.5 or ≤ −1.5 and q-value ≤ 0.05 are considered significantly differentially expressed. (H) Top 10 gene ontology (GO) enrichment analysis of upregulated genes identifying biological processes associated with myelinating SCs compared to immature and repair SCs. (I) Top 10 GO enrichment analysis of upregulated genes identifying biological processes associated with repair SCs compared to myelinating SCs.

During differentiation into myelinating SCs, differential expression analysis identified 726 upregulated and 947 downregulated genes compared to immature SCs (Figure 2E, full data in **Supplementary Table 1**). Myelinating SCs exhibited significant upregulation of KROX20/Egr2 and MBP genes, both essential for myelin formation. Gene ontology (GO) enrichment analysis identified the top 10 biological pathways enriched in myelinating SCs, including Eph/ephrin signaling, myelination, receptor tyrosine kinase signaling, neuron projection development, and sodium/potassium ion transmembrane transport (Figure 2H, full data in **Supplementary Table 2**). Gene set enrichment analysis (GSEA) corroborated enrichment in pathways associated with myelination, myelin maintenance, and membrane potential regulation, indicating the specialized role of myelinating SCs in axon wrapping, myelin sheath formation, and signal propagation (**Supplementary** Figure 2A).

Conversely, reprogramming to repair SCs was characterized by 430 upregulated and 380 downregulated genes compared to the myelinating stage (Figure 2F, full data in **Supplementary Table 3**). This transition was marked by downregulation of myelin-related genes, including MBP and KROX20/Egr2, and upregulation of transcription factors c-Jun, SOX2, and neurotrophin receptor p75^NTR^/Ngfr, indicating activation of a proliferative state (Figure 2F). GO analysis identified the enriched pathways involved in axon guidance, which facilitates axon regrowth and SC migration, as well as PI3K/Akt signaling, ERK1/2 cascade, and MAPK activation, which promote cell survival and regeneration. Additional enriched pathways included negative regulation of apoptosis, essential for preventing neuronal death, and positive regulation of cell migration, which directs SCs to injury sites (Figure 2I, full data in **Supplementary Table 4**). GSEA identified gene sets associated with neurogenesis, neuroinflammatory responses, growth activity, and apoptosis regulation, indicating their roles in nerve repair, inflammation response, and neuroprotection (**Supplementary** Figure 2B). Collectively, these transcriptomic data delineate stage-specific regulatory mechanisms that direct the functional specialization of SCs in myelinating and repair stages.

### 3. Isolation and characterization of extracellular vesicles from Schwann cells across distinct differentiation stages

To elucidate the role of SC-EVs in nerve repair, EVs were isolated from the conditioned media of immature, myelinating, and repair-stage SCs using a dielectrophoresis-based microfluidic device (iDEP) (Figure 3A), which we have previously established (34–36). The presence of EVs was validated according to the Minimum Information for Studies of Extracellular Vesicles (MISEV2023) guidelines (37). Nanoparticle tracking analysis (NTA) revealed a consistent mean diameter less than 200 nm of SC-EVs from all differentiation stages, characteristic of small EVs (Figure 3B) (37). Western blotting confirmed the presence of common EV markers including CD63, HSP70, and TSG101, and the absence of negative marker Calnexin (Figure 3C, the full-length western blot images were available in **Supplementary** Figure 3). These results support the integrity and specificity of EV preparations for further analyses.

**Figure 3.**
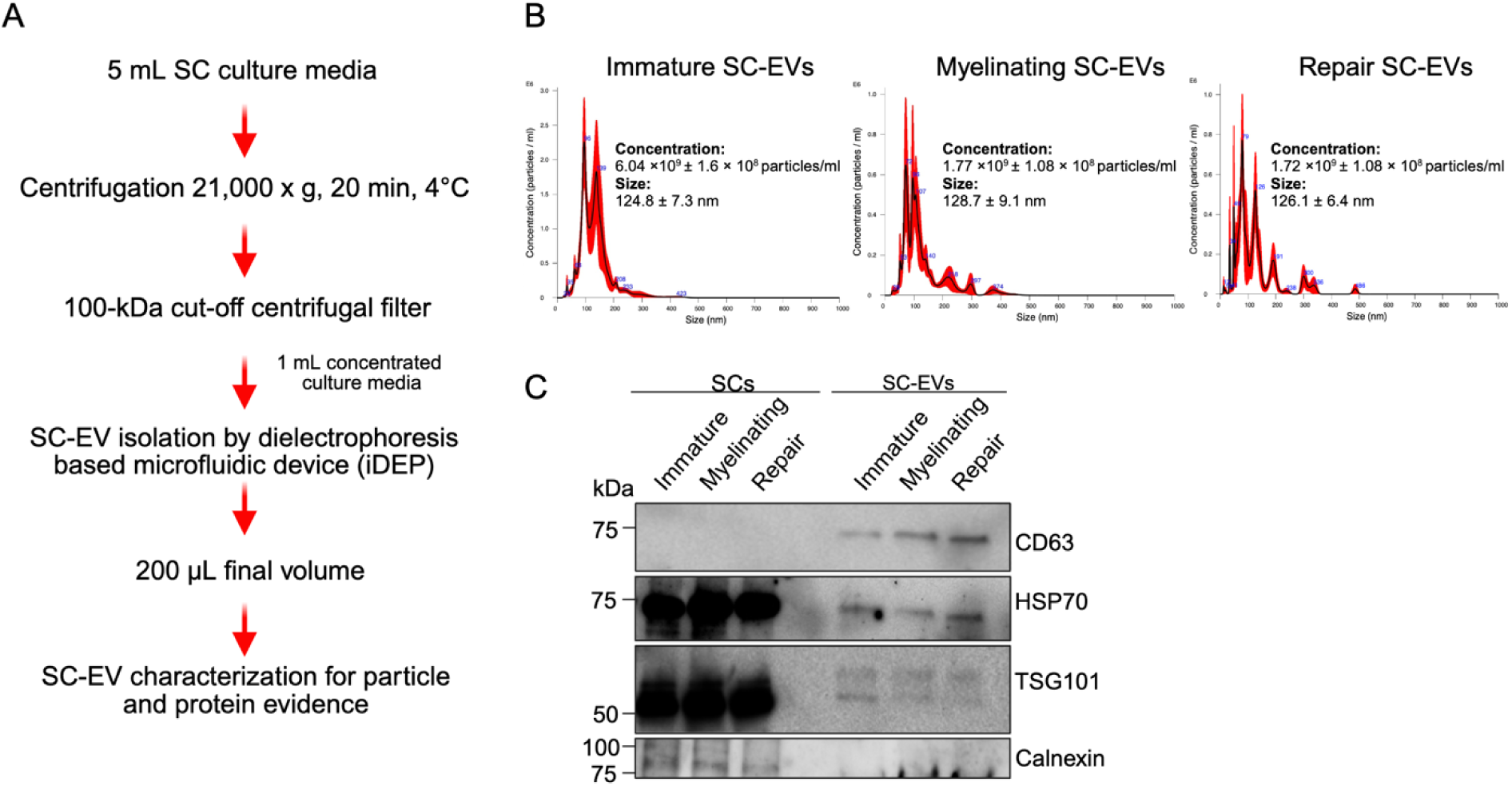
SC-EV isolation and characterization. (A) A schematic diagram represents SC-EV isolation process. (B) Nanoparticle tracking analysis (NTA) showing EV size distributions and concentrations, with average diameters of 124.8 ± 7.3 nm, 128.7 ± 9.1 nm, and 126.1 ± 6.4 nm, and concentrations of 6.04 × 10⁹, 1.77 × 10⁹, and 1.72 × 10⁹ particles/ml for immature, myelinating, and repair SCs, respectively. (C) Western blot analysis confirming the presence of EV-specific markers CD63, TSG101, HSP70, and calnexin across all stages.

### 4. Myelinating Schwann cell-derived extracellular vesicles carry regenerative proteins that may support neuroregeneration

To confirm the SC origin of EVs at different differentiation stages, super-resolution microscopy was employed to identify CD63⁺ vesicles co-expressing the SC marker p75^NTR^ (Figure 4A). Next, we utilized western blotting to further characterize their protein cargo. As expected, myelin basic protein (MBP) was highly enriched in myelinating SC-EVs, reflecting the identity of their parental cells (Figure 4B**, C**). Interestingly, SOX2 and p75^NTR^, markers of proliferation and repair, were enriched in myelinating SC-EVs despite relatively low expression in myelinating SCs, while their levels were reduced in SC-EVs from immature and repair stages compared to their cellular sources (as shown in Figure 1E). These findings suggest that myelinating SC-derived EVs selectively package regenerative proteins, highlighting their potential role in SC-neuron communication during peripheral nerve repair.

**Figure 4.**
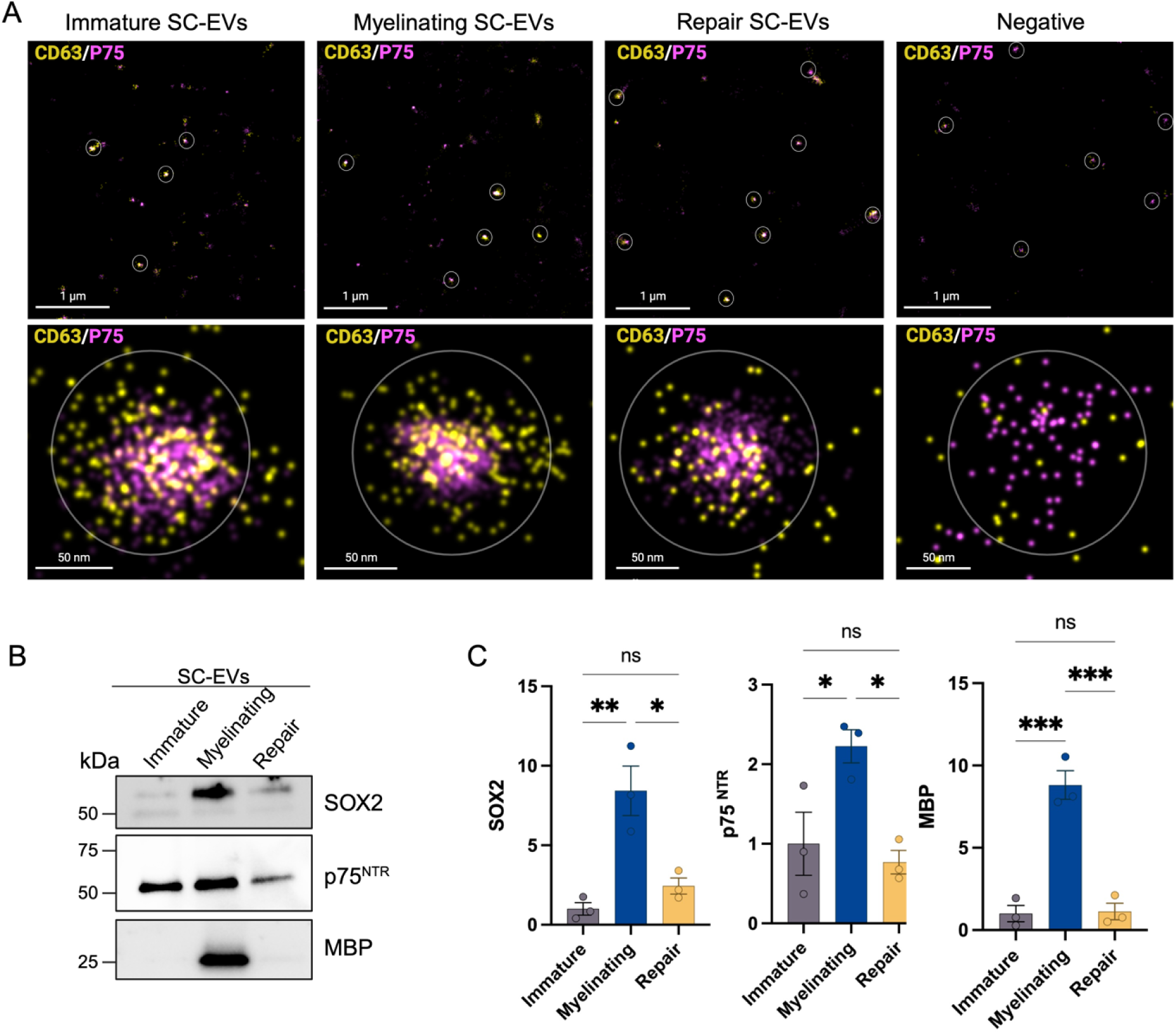
Stage-specific protein cargo in Schwann cell-derived extracellular vesicles (SC-EVs). (A) Representative direct stochastic optical reconstruction microscopy (dSTORM) showing positive CD63 signals co-expression with SC marker p75^NTR^ on a single particle SC-EV. Upper panels show lower magnification views (scale bar, 1 µm) with circled regions indicating EVs positive for both markers. Lower panels present high-magnification views of representative EVs (scale bar, 50 nm). (B) Western blot analysis of stage-specific EV marker SOX2, p75^NTR^, and MBP in EVs derived from immature, myelinating, and repair SCs (the full-length blot images are available in Supplementary figure 4). (C) Quantification of EV-associated protein expression demonstrating significant enrichment of SOX2, p75^NTR^, and MBP markers in EVs derived from myelinating SCs. Data presented as mean ± SEM (n = 3 biological replicates per group). Statistical analysis was performed by using one-way ANOVA with Tukey’s multiple comparisons test: * p < 0.05, ** p < 0.005, and *** p < 0.001, ns, not significant.

### 5. Stage-specific SC-EV miRNAs regulate pathways associated with nerve regeneration

To determine whether SC-EVs carry stage-specific miRNAs associated with nerve regeneration, we performed small RNA sequencing on EVs isolated from SC at three differentiation stages. Heatmap with hierarchical clustering of significantly expressed miRNAs revealed stage-specific expression profiles (Figure 5A), indicating selective miRNA cargo associated with each SC stage. We selected differentially expressed miRNAs from this heatmap, focusing on those upregulated or downregulated in repair SC-EVs compared to myelinating SC-EVs, as this transition represents a shift from axonal maintenance to regenerative function. In total, 52 downregulated and 67 upregulated mature miRNAs were identified in repair SC-EVs (**Supplementary Table 5**). Target prediction using the intersection between miRWalk and miRDB revealed 2,541 shared target genes of downregulated miRNAs and 2,325 targets of upregulated miRNAs (Figures 5B**, 5F**).

**Figure 5.**
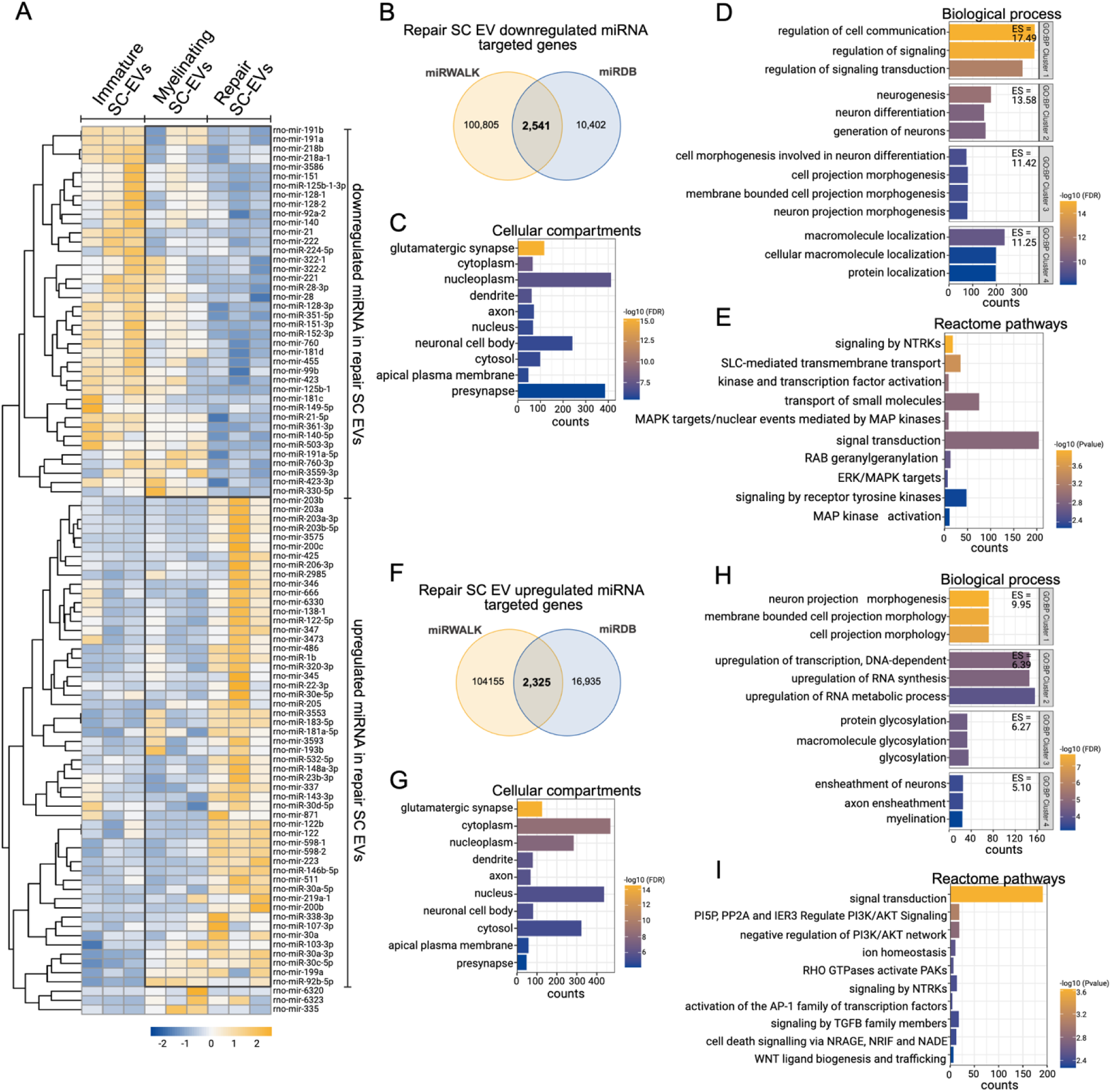
miRNA profiling of Schwann cell-derived extracellular vesicles (SC-EVs). (A) Heatmap with hierarchical clustering shows differentially expressed miRNAs in SC-EVs from immature, myelinating, and repair SCs. Each row represents an individual miRNA, with color intensity indicating relative expression levels (blue: downregulated; yellow: upregulated). (B, F) Venn diagrams showing the overlap of predicted target genes for downregulated (B) and upregulated (F) miRNAs in repair SC-EVs based on miRWalk and miRDB databases. The intersection represents commonly predicted target genes. Gene Ontology (GO) and Reactome pathway enrichment analysis of target genes corresponding to downregulated (C-E) and upregulated (G-I) miRNA in repair SC-EVs. Enrichments are categorized by cellular components (C, G), biological process clusters (D, H), and reactome pathways (E, I) to illustrate functional pathway involvement.

To assess the functional relevance of these miRNAs, we performed pathway enrichment analysis using DAVID. Gene ontology (GO) analysis of target genes of downregulated miRNAs in repair SC-EVs revealed significant enrichments in neuronal structures including dendrites, axons, neuronal cell bodies, and synapses (Figure 5C, full data in **Supplementary Table 6**) as well as biological processes involved in regulation of signaling (GO: BP cluster 1), neurogenesis (GO: BP cluster 2), neuron morphogenesis (GO: BP cluster 3) and protein localization (GO: BP cluster 4) (Figure 5D, full data in **Supplementary Table 7**). Reactome pathway enrichment showed involvement in signal transduction, MAPK signaling (including ERK/MAPK targets and MAP kinase activation), receptor tyrosine kinase signaling, and transmembrane transport (Figure 5E, full data in **Supplementary Table 8**).

Upregulated miRNAs in repair SC-EVs were also linked to target genes; presumably suppressed in recipient cells, that are involved in neuronal cell bodies, synapses, and dendritic structures (Figure 5G, full data in **Supplementary Table 6**), while the enriched biological processes were related to neuron projection morphogenesis (GO: BP cluster 1), transcriptional activation (GO: BP cluster 2), glycosylation (GO: BP cluster 3), and myelination (GO: BP cluster 4). (Figure 5H, full data in **Supplementary Table 7**). Reactome pathway analysis revealed significant enrichment in signaling pathways related to PI3K/AKT signaling, RHO GTPase pathways, TGFB and WNT signaling, transcription factor activation (including AP-1), ion homeostasis, and cell death signaling (Figure 5I, full data in **Supplementary Table 8**). Together, these results demonstrate that SC-EVs exhibit stage-specific miRNA cargo capable of modulating gene expression in recipient cells, thereby contributing to distinct roles in peripheral nerve regeneration.

### 6. Repair SC-EVs contain lncRNAs that may modulate miRNA activity

Long non-coding RNAs (lncRNAs) regulate gene expression through diverse mechanisms, including by acting as competitive endogenous RNAs or sponges that sequester miRNAs and modulate downstream targets (38, 39). To explore the presence of such regulatory lncRNAs in SC-EVs, we perform expression profiling across immature, myelinating, and repair SC-EVs. A heatmap of the most abundantly expressed lncRNAs in repair SC-EVs is shown in Figure 6A (expression data in **Supplementary Table 9**); although these lncRNAs were not statistically differentially expressed, they were selected based on relative abundance and remain uncharacterized in current databases. To assess their potential function as miRNA sponges, we selected upregulated clusters from the heatmap and performed miRNA-lncRNA interaction analysis using miRanda, focusing on high-affinity binding pairs (total score >140; minimum energy < –20 kcal/mol) (40). This analysis targeted on miRNAs downregulated in repair SC-EVs that are associated with neurogenesis. Upregulated lncRNAs were predicted to bind these miRNAs (Figure 6B, full data in **Supplementary Table 10**). These findings suggest that repair SC-EVs may deliver lncRNAs sequester miRNA-mediated repression of pro-regenerative genes, thereby enhancing their neuroregenerative potential.

**Figure 6.**
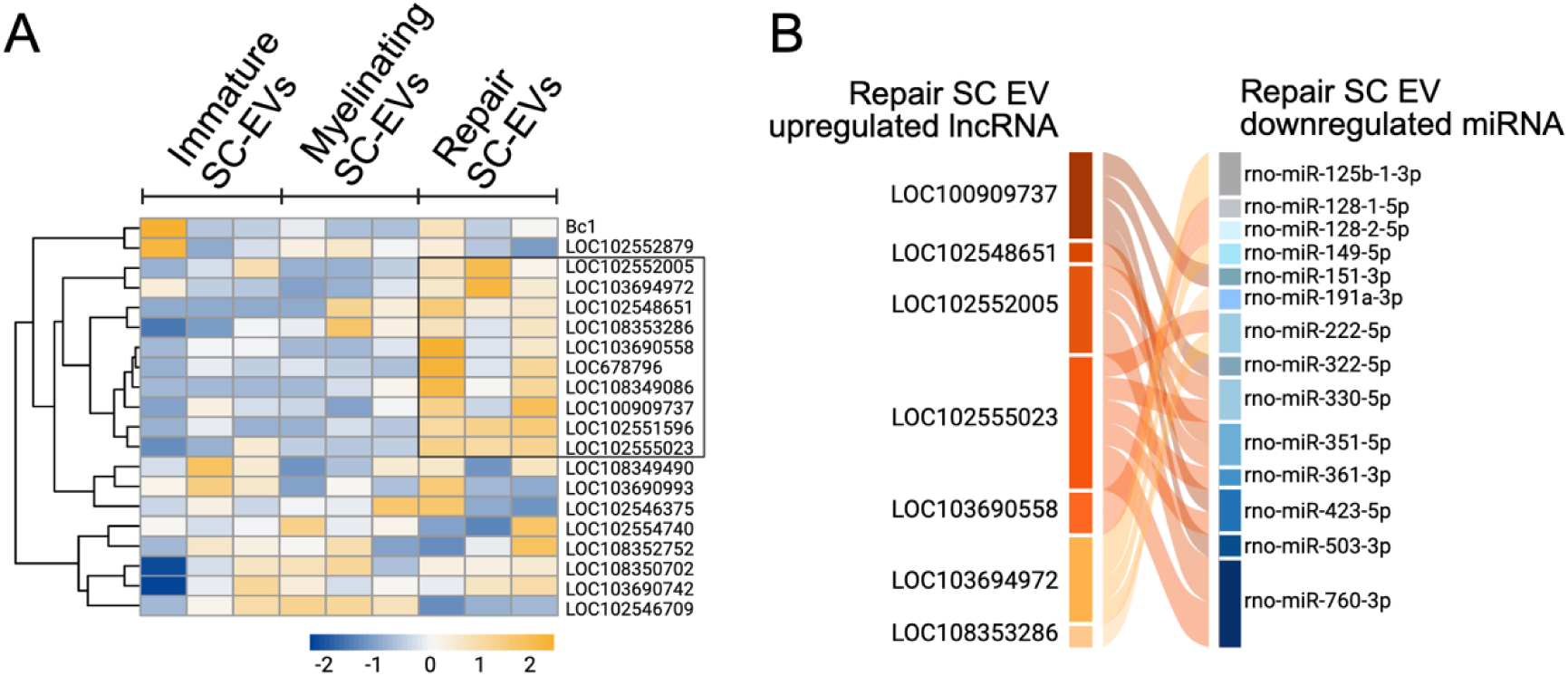
lncRNA expression and predicted miRNA interactions in repair SC-EVs. (A) Heatmap with hierarchical clustering of the most abundant lncRNAs in repair SC-EVs. Each row represents an individual lncRNA, with color intensity indicating relative expression (blue: low; yellow: high). (B) Sankey diagram showing predicted interactions between upregulated lncRNAs in repair SC-EVs and downregulated miRNAs, based on miRanda binding predictions.

## Discussion

Following nerve injury, SCs undergo a dynamical phenotypic reprogramming, transitioning from a myelinating to a repair stage to support nerve regeneration. Emerging evidence suggests that SC-EVs are crucial mediators of SC-neuron communication (14, 23–25). However, the molecular cargo and functional divergence of SC-EVs at distinct differentiation stages remain incomplete understood. In this study, we comprehensively profiled proteins, miRNAs, and lncRNA cargo of SC-EVs of immature, myelinating and repair phenotypes. Our results reveal stage-specific regulatory signatures that suggest distinct contributions in peripheral nerve regeneration.

To establish a robust experimental model, we first validated the differentiation stages of immature, myelinating and repair SCs (31) by confirming characteristic morphology and molecular markers (Figure 1). Immature SCs showed high expression of c-Jun, SOX2, and p75^NTR^; genes associated with proliferation, developmental plasticity and regenerative potential (31, 32, 41, 42). Myelination induced upregulation of KROX20 and MBP and morphological changes characteristic of mature myelinating SCs (32, 41, 42). Upon dedifferentiation, repair SCs re-acquired immature-like morphology and re-expressed regenerative markers, highlighting SC plasticity in response to injury-associated signals (13, 31, 33, 43, 44). Transcriptomic analyses further supported these phenotypic transitions. Myelinating SCs exhibited elevated expression of myelin-related genes, including KROX20/Egr2 and MBP, and activation of signaling pathways critical for myelination, such as receptor tyrosine kinase signaling and Eph/ephrin signaling. Upregulation of ion channel-related genes further reflected the functional maturation of myelinating SCs. (Figure 2H) (1, 13, 45). In contrast, repair SCs showed enriched signaling pathways linked to proliferation, axon guidance, anti-apoptosis, and immunomodulation (Figure 2I), consistent with their role in supporting regeneration (7, 46). This was accompanied by reactivation of c-Jun, SOX2, and p75^NTR^, and downregulation of myelin-specific genes; the hallmarks of repair phenotype.

SC-EVs carry stage-specific molecular cargo that mirrors the phenotype and regenerative capacity of their cells of origin. Notably, EVs released by myelinating SCs were enriched in regenerative proteins, including SOX2 and p75^NTR^, despite low intracellular expression of these factors at this stage (Figure 1E**, 4B**). This discrepancy points to selective cargo-sorting mechanisms, whereby EV content does not directly reflect cellular protein abundance (47, 48). One possibility is that SCs utilize EVs to expel proteins no longer required intracellularly, consistent with the dual roles of EVs in both intercellular communication and cellular waste management (49, 50). The enrichment of SOX2 and p75^NTR^ within myelinating SC-EVs suggests their functional role in nerve repair. Since myelinating SCs mostly present near injury sites, their EVs may serve as readily accessible sources of regenerative signals during the early phases of nerve regeneration. This mechanism is supported by previous studies demonstrating that SOX2-contained EVs promote neurite extension and axonal regeneration (26, 51), and p75^NTR^ enhances neuronal survival (52). Nevertheless, functional validation is required to confirm whether EV-contained SOX2 and p75^NTR^ actively contribute to nerve regeneration in this context.

A particular novel aspect of our study is the identification and functional annotation of miRNAs selectively packaged into SC-EVs from repair SCs that were dedifferentiated from a myelinating stage. Small RNA sequencing of SC-EVs from three differentiation stages revealed stage-specific miRNA profiles, with repair SC-EVs showing distinct expression patterns compared to immature and myelinating SC-EVs (Figure 5A). We focused on differentially expressed miRNAs between repair and myelinating SC-EVs, as this transition represents a key shift from axonal maintenance to regenerative activation. Notably, repair SC-EVs exhibited distinct miRNA signatures predicted to target neuronal compartments (Figure 5C**, 5G**), suggesting that these EV-associated miRNAs modulate genes implicated in synaptic structure and neuronal architecture. Many of upregulated miRNAs were predicted to repress transcripts involved in neuron projection morphogenesis, RNA biosynthesis, protein glycosylation, and myelination (Figure 5H). This repression likely transiently limits transcriptional activity and structural differentiation, consistent with the early repair phase’s prioritization of axon outgrowth and extension over maturation and synaptic stabilization (23, 26). Pathway analysis supported this regulatory role, revealing predicted suppression of PI3K/AKT, NTRK, RHO GTPase, and AP-1 signaling pathways (Figure 5I), which typically promote neuronal differentiation, axon elongation and synaptic formation (53–55). In parallel, repression of genes associated with axon ensheathment suggests that repair SC-EVs may delay myelination in both neurons and SCs, promoting a pro-repair phenotype (56). Additionally, suppression of cell death signaling supports a neuroprotective function during the early injury response (57). In contrast, downregulated miRNAs in repair SC-EVs were associated with the de-repression of regeneration-associated genes silenced under homeostatic conditions, including those involved in regulation of signaling, neurogenesis, cell projection morphogenesis, macromolecular transport, and protein localization (Figure 5D). This de-repression may promote neurite outgrowth, cytoskeletal remodeling, and intercellular communication, supporting structural reformation of injured axons and dendrites. Reactome pathway analysis further highlighted activation of NTRK signaling, MAPK cascade, and receptor tyrosine kinase pathways (Figure 5E), which are essential for neurotrophin-mediated axon elongation and neuronal survival (54). The miRNA shift observed may be related to the differential expression of p75^NTR^, which is elevated in repair SCs and reduced in myelinating SCs. Recent work demonstrated that genetic deletion of p75^NTR^ and its co-receptor sortilin in SCs significantly altered EV-associated small RNA profiles, including miRNAs involved in autophagy and phosphatidylinositol signaling (58). These findings reveal that SC-EVs act as dynamic regulators of gene expression during nerve injury, with their miRNA cargo contributing to both intra- and intercellular reprogramming. This is consistent with previous studies showing that SC-EVs promote neurite outgrowth, axonal regrowth, and modulation of regenerative microenvironments (26, 29). Additionally, our transcriptomic data further support these regulatory roles, revealing that repair SCs upregulate genes involved in axon regrowth, migration, and modulation of PI3K/Akt, ERK, and MAPK pathways, consistent with the predicted functions of EV miRNAs and suggesting that SC themselves may also be targets of EV-mediated signaling.

Among miRNAs identified in SC-EVs, several have established roles in promoting nerve regeneration. For example, miR-125b supports neurite elongation (59), miR-138-5p inhibit apoptosis (60), and miR-193b-3p and miR-223-3p contribute to anti-inflammation (61, 62). Similarly, miR-345-3p implicates in resolving inflammation and promotes neuronal survival (63). Conversely, downregulation of miR-125b, miR-140, miR-181c, and miR-199a-3p has been linked to enhanced axonal extension and cell viability (64–67), suggesting that suppression of specific miRNAs may de-repress neuronal regeneration. While miR-21 is widely recognized as pro-regenerative (26, 68), its downregulation in our dataset may reflect dynamic or stage-specific regulation during the repair process (69). Furthermore, suppression of miR-423-5p and miR-92a-2-5p, both involved with neuroprotection and neuronal differentiation (70, 71), are consistent with transcriptional reprogramming toward a regenerative stage. Notably, several miRNAs in SC-EVs remain uncharacterized, presenting new opportunities to uncover regulatory elements underlying nerve regeneration.

In parallel, we identified highly expressed long non-coding RNAs (lncRNAs) enriched in repair SC-EVs (Figure 6), most of which are currently unannotated. Given that lncRNAs can function as ceRNAs or molecular sponges that sequester miRNAs and relieve suppression of their target transcripts (38, 39), we assessed potential lncRNA–miRNA interactions using miRanda (40). lncRNAs exhibited strong predicted binding to miRNAs downregulated in repair SC-EVs, i.e., miR-125b, miR-149-5p, miR-222-5p, miR-330, miR-423-5p, and miR-128-3p, all of which are implicated in neuroregenerative processes (64, 70, 72–74). These interactions demonstrated low hybridization energies and multiple binding sites, indicating stable lnc-RNA-miRNA complexes (**Supplementary Table 10**). Together, these findings support a model in which lncRNAs within SC-EVs post-transcriptional regulation by modulating miRNA activity during nerve injury. Future studies will be essential to validate these lncRNA–miRNA interactions and elucidate their contribution to SC plasticity and peripheral nerve regeneration.

This study establishes a transcriptomic framework for understanding the stage-specific molecular cargo of SC-EVs and their potential regulatory roles in nerve regeneration. However, one limitation is the potential carryover of EVs generated during earlier differentiation stages but not yet released from multivesicular bodies, which may confound the interpretation of stage-specific cargo profiles. ManNAz metabolic labeling combined with click chemistry offers a promising strategy to timestamp newly synthesized EVs, potentially enhancing the precision of stage-specific EV profiling in future studies (75). Our omics-based analyses highlight candidate mechanisms, including lncRNA-mediated miRNA modulation, but these remain predictive. lncRNAs may also function through other mechanisms such as chromatin remodeling, transcriptional regulation, and modulation of RNA-binding proteins (38), which warrant future investigation. A major limitation of this study is the lack of functional validation to confirm the biological relevance of the identified molecular signatures, particularly top-ranked miRNAs and lncRNAs. Future studies employing multicellular systems, three-dimensional culture models, and *in vivo* approaches will be essential. Moreover, engineering strategies to enhance SC-EV production and regenerative potency represent promising avenues for translational development. Notably, mechanical and electrical stimulation of SCs has been shown to increase EV secretion and enrich their cargo with regenerative factors (14, 29). Incorporating piezoelectric or conductive biomaterials (76–79) into engineered culture systems may further enable stage-specific EV enrichment. Together, these approaches lay the groundwork for advancing SC-EVs toward translational SC-EV therapies for peripheral nerve repair.

## Materials and Methods

### Rat primary Schwann cell culture

Primary Rat Schwann cells (RSC) (Catalog R842-05a) were purchased from CELL Applications (San Diego, CA, USA) and maintained according to the manufacturer’s protocol. Briefly, Schwann cells were cultured in RSC growth medium (Catalog R825-500) (CELL Applications, San Diego, CA, USA) at 37°C in 5% CO_2_. The culture medium was replaced every 48 h. Sub-culturing was performed when the cells reached 80% confluency and cells were maintained for no longer than 5 passages. Passages 3 to 5 were used for experiments.

### Cell differentiation and dedifferentiation assay

Six-well plates and 11 mm glass coverslips were incubated with Schwann Cells Coating solution (Cat036-20) CELL Applications (San Diego, CA, USA) at 37°C overnight. The remaining solution was aspirated, rinsed twice with PBS, and dried. RSC samples were prepared according to a modified protocol (31). Cells were seeded into a coated 6-well plate at a density of 2×10^5^ cells/well for western blot and RNA sequencing and seeded into a 24-well plate (covered with coated glass coverslips) at a density of 1×10^4^ cells/well for immunofluorescence staining. Cells were cultured in RSC complete growth medium (R825-500) for 24 h and the medium was changed into the complete medium-RSC basal media containing 10% Exosome-Depleted FBS (Fisher), 2mM of Foskolin (Sigma-Aldrich, catalog number: F6886), 10 ng/mL of human neuregulin 1-β1/heregulin (R&D Systems, Catalog Number: 396-HB/CF), and 100 mg/mL penicillin-streptomycin (Gibco) for 24 h. Then the medium was removed and replaced by a medium containing DMEM, 1% Exosome-depleted FBS, and 100 mg/mL penicillin-streptomycin (starvation medium) overnight to limit mitogenic stimulation. Cells were then differentiated into myelinating SC by removing the starvation medium and replacing it with medium containing DMEM, 1% Exosome-depleted FBS, 100 mg/mL penicillin-streptomycin, and 1 mM db-cAMP (Sigma, USA). After 72 h, the medium was replaced with the Exosome-depleted complete medium for an additional 72 h to induce dedifferentiation. Cells maintained in the absence of db-cAMP-inducing agents served as a control for undifferentiated cells. Cells were monitored daily to observe changes in morphology. At the end of the assay, cells were harvested for SC stage-specific markers for characterization of Schwann cell differentiation stages, and the supernatant was collected for EV isolation.

### Immunofluorescence staining

Cells were washed three times before fixation in 3.7% formaldehyde for 15 min at room temperature. Following two washes with PBS, cells were permeabilized with 0.1% Triton X-100 (Fisher) in PBS at 4°C for 5 min and washed with PBS. The cells were blocked with 200uL 0.2% bovine serum albumin (BSA) in PBS (blocking buffer) for 1 h at RT, and then incubated with primary antibodies (1:400 diluted in blocking buffer) as follows; rabbit anti-p75 NGF Receptor (ab52987; Abcam, Waltham, MA, USA), rabbit anti-SOX2 (ab97959; Abcam, Waltham, MA, USA), mouse anti-c Jun (NBP2-71059, Novus Biologicals, CO, USA), rabbit anti-KROX20 (Abcam, Waltham, MA, USA) or EGR2 (ab245228; Abcam, Waltham, MA, USA) antibodies at 4°C overnight. The next day, cells were washed with PBS followed by separate incubations for 30 min at 37 °C with 1:100 dilution of rhodamine phalloidin (Invitrogen, OR, UAS), goat anti-rabbit IgG Alexa Fluor 488 (ab150077; Abcam, Waltham, MA, USA), and anti-mouse IgG Alexa Fluor 647 (ab150115; Abcam Waltham, MA, USA) secondary antibodies. After washing, cells were mounted in mounting medium with DAPI (Catalog-50011, ibidi, WI, USA) on a microscope slide. Clear nail polish was used to seal coverslips. Imaging was performed using a NIKON ECIPSE TE-2000-S.

### Schwann cell-derived extracellular vesicle isolation

EVs were isolated using an insulator-based dielectrophoretic (iDEP) as described (34, 80). The culture supernatant (5 mL) from each condition was centrifuged at 21,000×g at 4 °C for 20 minutes to remove large EVs and cell debris. Conditioned media were concentrated 5-fold using Amicon Ultra-4 centrifugal filters (100-kDa cutoff; Millipore, Burlington, MA, USA), which also removed a substantial portion of soluble proteins prior submitted to the iDEP. Briefly, borosilicate glass capillaries (BF-100-50-15, Sutter Instrument, Novato, CA, USA) were fabricated into micropipettes with 2 μm pore diameters using a laser-assisted puller P2000 (Sutter Instrument Company, Novato, CA, USA). The micropipette was filled with 0.02-μm filtered 1x PBS buffer using a 33-gauge Hamilton syringe needle and positioned on a substrate. Then, 1X filtered PBS and 50 μL of sample were loaded at the base side and tip side of the micropipette, respectively. EVs were trapped at the tip by applying a 10 V/cm direct current (DC) for 15 min, followed by a release in 15 μL 1× filtered PBS by reversing the applied voltage for another 10 min. Isolated EVs were aliquoted and stored at −80°C for further analysis. Only one freeze-thaw cycle was allowed for each aliquot.

### Western Blot Analysis

Cell and EV samples were lysed in 1X RIPA buffer (Abcam) and 1X protease/phosphatase inhibitor (Abcam). The samples were subjected to measure protein concentration using the DC Protein Assay (Bio-Rad, Hercules, CA, USA). The cell lysates (5ug) and EV lysates (1ug) were mixed with 4x Laemmli buffer (Bio-Rad), heated at 95°C for 5 mins. The samples were run into a 4–20% Mini-PROTEAN TGX Precast Protein gel, 200V for 35 min before being transferred onto a PVDF membrane using a Trans-Blot Turbo System (Bio-Rad). The membrane was blocked with EveryBlot blocking buffer (Biorad) for 5 min, and then probed with 1:1,000 dilution of primary antibodies as follows; rabbit anti-p75 NGF Receptor (ab52987; Abcam, Waltham, MA, USA), rabbit anti-SOX2 (ab97959; Abcam), rabbit anti-c Jun (ab40766; Abcam), rabbit anti-KROX20 or EGR2 (ab245228; Abcam) and human anti-myelin basic protein (ab209328; Abcam) antibodies for SC markers, mouse anti-beta actin antibody (Ab6276; Abcam), and mouse anti-CD63 (Ab108950; Abcam), rabbit anti-TSG101 (ab125011;Abcam) and rabbit anti-HSP70 (ab181606; Abcam) antibodies for EV common markers at 4°C overnight. After washing, the membranes were incubated with secondary antibody specific to the species of the primary antibody at a dilution of 1:2000 as followes; goat anti-rabbit IgG-HRP, goat anti-mouse IgG-HRP (Abcam), and goat anti-human IgG-HRP (Abcam) at room temperature for 1 h. The immunoblot was developed by ECL (Bio-Rad, USA) for the cell samples and SuperSignal West Pico PLUS (Thermo) for the EV samples. The membrane was imaged on a ChemiDoc MP imaging system (Bio-Rad, USA). Signal intensities were quantified using the Image J software (NIH, USA). At least three independent experiments were performed. For SCs, protein bands were normalized to actin and quantified relative to the control (undifferentiated condition). For EVs, band intensities were quantified relative to the control (undifferentiated condition).

### Nanoparticle tracking analysis (NTA)

A NanoSight NS300 (Malvern Instruments Ltd., Malvern, Worcestershire, UK), integrated with a sample pump, was utilized for the analysis. EV samples were diluted in 0.22 μm filtered 1x PBS at a ratio of 1:40 to achieve a total volume of 1 ml. The diluted sample was then injected into the sample chamber using the syringe pump. Five 1-minute videos were recorded under the following parameters: camera: sCMOS; cell temperature: 25°C; syringe pump speed: 30 μl/s. After capture, the videos were analysed by NanoSight Software NTA 3.4 Build 3.4.003 using the settings: detection threshold, 5; blur size and max jump distance, auto. Ideal concentrations contained 20–100 particles/frame.

### Super-resolution microscopy

Single-level EV images were obtained using a Nano imager S Mark II microscope (Oxford Nanoimaging, Oxford, UK) equipped with a 100X, 1.4 NA oil immersion objective, an XYZ closed-loop piezo 736 stage, and dual or triple emission channels split at 640 and 555 nm. EV samples were prepared according to a modified protocol (81). 1×10^9^ EVs were stained with 5ug of primary antibodies in combination of mouse anti-CD63 (Ab108950; abcam) and rabbit anti-p75 NGF Receptor (ab52987; abcam) in 100 μL 0.2% BSA in PBS overnight on a shaker at 4 °C. Next day, samples were wash with 400 μL filtered 1xPBS for three times using Ultracel-100 K membrane filter (Merck Millipore, Burlington, MA, USA) at 5000 RPM for 3 min. Samples were collected and labeled with secondary antibodies including Goat pAb to mouse IgG AF488 (AB150116) and Goat pAb to rabbit IgG AF594 (Abcam, Waltham, MA, USA) at a dilution of 1:400 in 400 μL 0.2% BSA in PBS for 3h at 4 °C. Following reaction completion, samples were washed three times with filtered 1X PBS and collected in a final volume of 100 μL. For imaging, μ-Slide 8 well glass bottom slides (Ibidi, Fitchburg, WI, USA) were washed two times with distilled water, coated with 100 uL poly-L-lysine solution (Sigma Aldrich, St. Louis, MO, USA), and incubated at 37°C overnight. Upon incubation, the solution was carefully removed and the sample of antibodies-EVs was added at a final volume of 100 μL and kept at 4 °C overnight. The next day, the buffer was removed, washed, and fixed with 3.7% formaldehyde for 15 minutes at 4°C. Lastly, samples were washed 3X with filtered PBS. Three to five fields of view were recorded for each sample using direct stochastical optical reconstruction microscopy (dSTORM). Analysis was performed using algorithms including filtering, drift correction, and DBScan clustering developed by ONI (Oxford Nanoimaging, Oxford, UK) via the Collaborative Discovery (CODI) platform.

### Whole RNA Sequencing for mRNA and lncRNA

Whole-transcriptome RNA sequencing was performed by the Genomics, Epigenomics, and Sequencing Core (GESC) at the University of Cincinnati following updated protocols as previously described (82, 83). Total RNA was isolated from Schwann cells for mRNA-seq and SC-EVs for lncRNA-seq, and quality control was assessed using the Agilent Bioanalyzer with the RNA 6000 Pico Kit (Agilent Technologies, Santa Clara, CA). Next, 10 ng of total RNA was used as input for library preparation using the NEBNext Ultra II Directional RNA Library Prep Kit (New England BioLabs, Ipswich, MA). PCR was run for 9 cycles. Library quality and concentration were evaluated using Qubit fluorometric quantification (Thermo Fisher Scientific, Waltham, MA). Each library was then uniquely indexed and proportionally pooled. The final library pool was sequenced on the Illumina NextSeq 2000 platform (Illumina, San Diego, CA) with setting of PE 2×61 bp to generate ∼44 million reads. Sequencing output was processed via the Illumina BaseSpace Sequence Hub to generate fastq files for downstream mRNA and long noncoding RNA (lncRNA) analysis.

### MicroRNA-Sequencing

Small RNA sequencing was carried out by the GESC following established protocols (84, 85). Total RNA isolated from SC-EVs was quantified and quality control-assessed using the RNA 6000 Pico Kit on the Agilent Bioanalyzer (Agilent Technologies, Santa Clara, CA). ∼5 ng of total RNA was used for library construction with the NEBNext Multiplex Small RNA Library Prep Kit (New England BioLabs, Ipswich, MA), incorporating a protocol modification to increase sensitivity and specificity for small RNA detection. PCR amplification was performed with 15 cycles, and 10 μL of each uniquely indexed PCR mixture was pooled. The pooled libraries were then subjected to DNA purification using the DNA Clean & Concentrator kit (Zymo Research, Irvine, CA). For precise size selection, the libraries were spiked with custom 135 and 146 bp DNA ladders and run on a high-resolution agarose gel electrophoresis. MiRNA-containing fragments, ranging from 135 to 146 bp (including adapters and miRNA inserts), were cut, gel-purified, and quantified using the NEBNext Library Quantification Kit on a QuantStudio 5 Real-Time PCR System (Thermo Scientific). Quantified libraries were low-depth sequenced on the Illumina NextSeq 2000 platform (Illumina, San Diego, CA) for the first round to generate a few million reads per sample. Based on the preliminary read counts, library input volumes were adjusted to normalize sample representation. A second round, high-depth sequencing run was subsequently performed to generate ∼4.5 million reads per sample for final data analysis.

### Bioinformatics and Data Analysis

#### mRNA analysis

RNA sequencing reads in FASTQ format were first subsampled, and strandedness was inferred using fq tool (https://github.com/stjude-rust-labs/fq). Quality control was performed with FastQC (http://www.bioinformatics.babraham.ac.uk/projects/fastqc), followed by adapter and quality trimming using Trim Galore (https://www.bioinformatics.babraham.ac.uk/projects/trim_galore) and Cutadapt (86). Trimmed reads were aligned to the rat reference genome using the STAR (87), with HISAT2 (88) as an alternative aligner. The alignments were sorted and indexed using SAMtools (89), and PCR duplicates were marked using Picard MarkDuplicates (http://broadinstitute.github.io/picard/). Transcript assembly and quantification were performed using StringTie (90). BEDTools (91) and bedGraphToBigWig were used to generate bigWig tracks. Intensive quality control of the RNA-seq data was carried out using tools including RSeQC (92), Qualimap (93), and dupRadar (94). Transcript quantification was performed using Salmon (95), and transcript counts were converted to gene counts using the tximport package (https://github.com/thelovelab/tximport). Differential gene expression analysis between sample groups was conducted using DESeq2 v1.26.0 (96). Hierarchical clustering heatmaps of differentially expressed genes (DEGs: log₂ fold change ≥ |0.58| and q-value ≤ 0.05) were generated. Genes with log₂ fold change ≥ 1.5 or ≤ −1.5 and q-value ≤ 0.05 were considered significantly differentially expressed and visualized in volcano plots. All upregulated genes were submitted to DAVID (http://david.ncifcrf.gov) for gene ontology (GO) enrichment analysis (97, 98) to identify the top 10 biological processes ranked by fold enrichment (FDR < 0.05). Additionally, genes ranked by log₂ fold change were submitted for gene set enrichment analysis (GSEA) using the GSEA tool (99). Gene sets upregulated in the “na_pos” phenotype with FDR < 25% were considered significant. All visualization were generated using SRplot (100).

#### miRNA analysis

miRNA sequencing data were processed using the nf-core/smrnaseq pipeline (version 2.3.1), which performs adapter trimming, quality control, alignment, and quantification of small RNA reads. The pipeline generated both raw and normalized gene count files, with normalization reported in counts per million. Differential expression analysis between sample groups was conducted using the DESeq2 package (v1.26.0) in R (96). A summary report of quality control metrics and pipeline outputs was generated using MultiQC (101). Significantly differentially expressed miRNAs were identified based on log₂FoldChange ≥ |0.58| and false discovery rate (FDR) ≤ 0.05. DEGs clustering heatmaps were generated using SRplot (100), and the two primary clusters representing upregulated and downregulated miRNAs in myelinating versus repair SC-EV groups were selected for downstream analysis. Target prediction for differentially expressed miRNAs was performed using the intersection of miRWalk (version 3.0, binding P-value ≥ 0.95) (102) and miRDB (prediction score >50) (103). Predicted target genes were submitted to DAVID (http://david.ncifcrf.gov) for gene ontology (GO) analysis to identify the top 10 cellular compartments. The DAVID gene functional classification tool (http://david.ncifcrf.gov) (104) was used to identify four functional clusters ranked by enrichment score (P-value ≤ 0.05). All visualizations were performed using SRplot (100).

#### lncRNA analysis

RNA sequencing reads were assessed for quality using FastQC (https://github.com/stjude-rust-labs/fq)and subsequently trimmed to remove adapter sequences using Trim Galore (https://www.bioinformatics.babraham.ac.uk/projects/trim_galore) and Cutadapt (86). The cleaned reads were aligned to the rat reference genome using the STAR aligner (87). Raw counts and lncRNA expression (Transcripts Per Million, TPM) were generated. The top 20 most highly expressed lncRNAs were used to generate heatmap. Upregulated clusters from the heatmap were subjected to miRNA-lncRNA interaction prediction using miRanda through SRplot (100). Interactions with high binding affinity (total score >140 and minimum free energy < –20 kcal/mol) were selected (40). All data visualization and plotting were performed using SRplot (100).

## Data Availability

The data that support the findings of this study have been deposited in the NCBI Gene Expression Omnibus (GEO) under accession number GSE300703 and are publicly available at: https://www.ncbi.nlm.nih.gov/geo/query/acc.cgi?acc=GSE300703.

## Acknowledgments

This study was made successful through the support and collaboration of several core facilities and individuals. RNA isolation and sequencing were performed by the Genomics, Epigenomics, and Sequencing Core (GESC) at the University of Cincinnati. We gratefully acknowledge Xiang Zhang and Sam Bell from GESC, and the team at Information Services for Research (IS4R), Cincinnati Children’s Hospital Medical Center: Aditi Paranje, Ronika De, and Ashley Kuenzi for their invaluable bioinformatics expertise and technical support, which significantly contributed to the success of this work. Graphical abstract and Figure 1A were created with BioRender.com.

## Ethics approval

Ethics approval not required.

## Author contributions

**Manju Sharma**: Methodology, Investigation, Writing – review & editing, **Supasek Kongsomros**: Conceptualization, Methodology, Investigation, Visualization, Writing – original draft, Writing – review & editing, **Maulee Sheth**: Methodology, Investigation, Writing – review & editing, **Somchai Chutipongtanate**: Investigation, Writing – review & editing, **Leyla Esfandiari**: Conceptualization, Writing – review & editing, Supervision, Funding acquisition. Co-first authors: Manju Sharma and Supasek Kongsomros contributed equally to this manuscript, and each has the right to list themselves first in author order on their CVs.

