## Supporting Information for "Stage-specific extracellular vesicle cargo from Schwann cells orchestrates peripheral nerve regeneration"

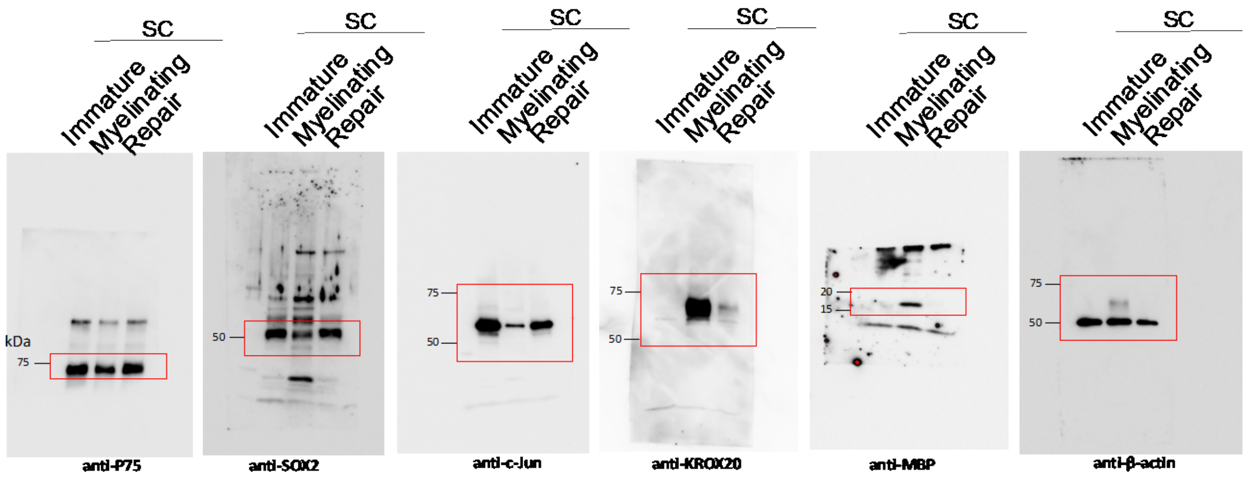


**Supplementary figure 1**. The full-length western blot results of the cropped images as presented in Figure 1E.

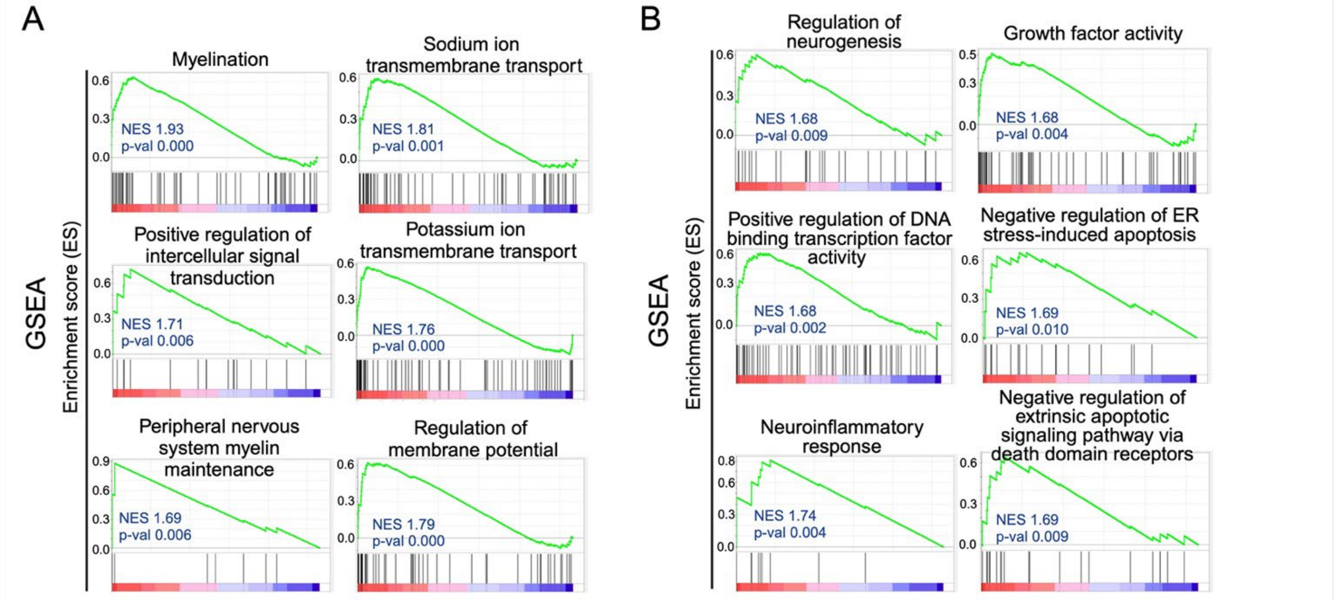


**Supplementary Figure 2** (A) positive gene set enrichment analysis (GSEA) of upregulated genes identifying biological processes associated with myelinating SCs compared to immature and repair SCs. (B) GSEA of upregulated genes identifying biological processes associated with repair compared to myelinating SCs.

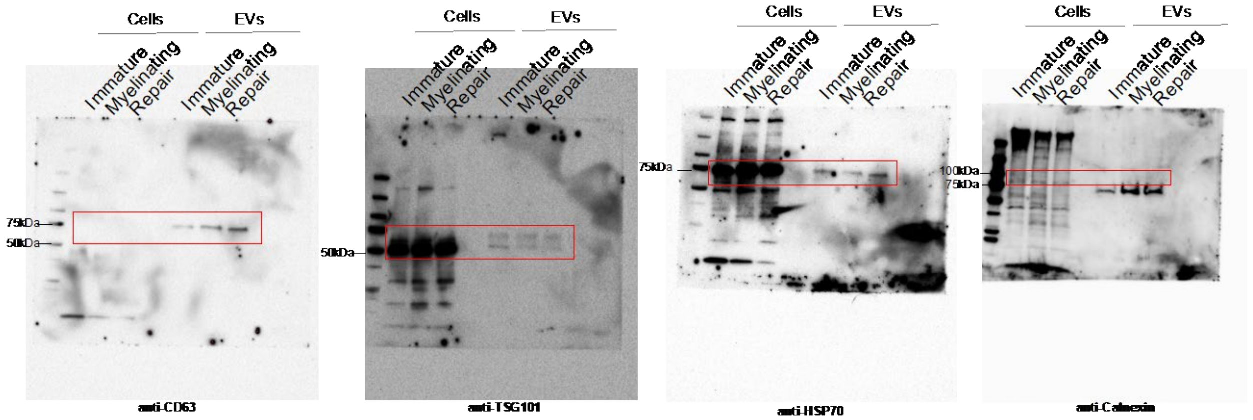


**Supplementary figure 3**. The full-length western blot results of the cropped images as presented in Figure 3C.

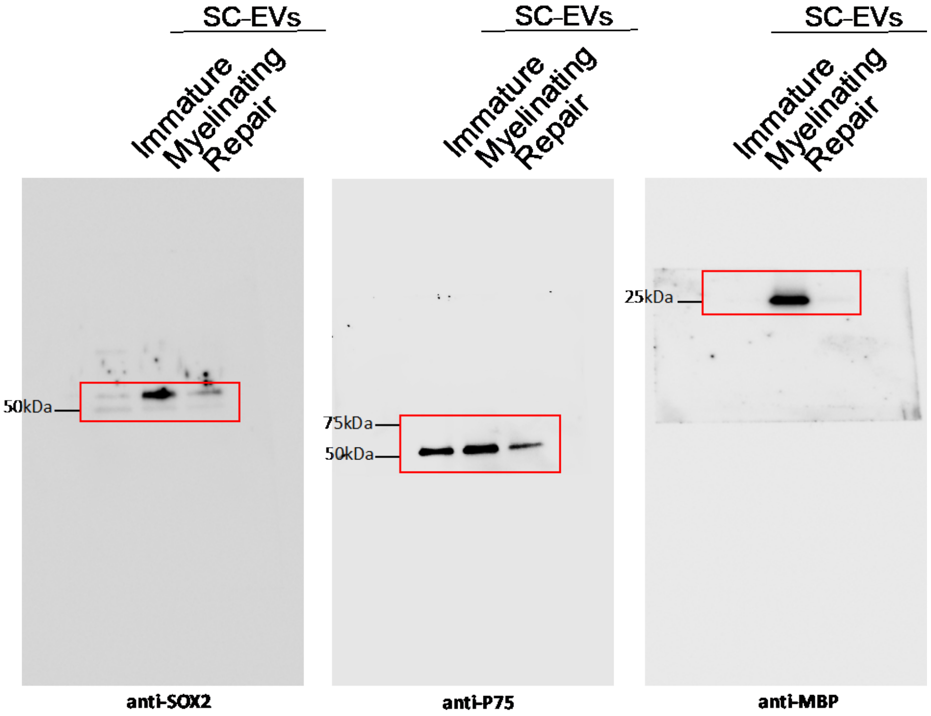


**Supplementary figure 4.** The full-length western blot results of the cropped images as presented in Figure 4B.
